# VIBES: A Workflow for Annotating and Visualizing Viral Sequences Integrated into Bacterial Genomes

**DOI:** 10.1101/2023.10.17.562434

**Authors:** Conner J. Copeland, Jack W. Roddy, Amelia K. Schmidt, Patrick R. Secor, Travis J. Wheeler

**Affiliations:** Division of Biological Sciences, University of Montana, Missoula, MT, USA; R. Ken Coit College of Pharmacy, University of Arizona, Tucson, AZ, USA

## Abstract

Bacteriophages are viruses that infect bacteria. Many bacteriophages integrate their genomes into the bacterial chromosome and become prophages. Prophages may substantially burden or benefit host bacteria fitness, acting in some cases as parasites and in others as mutualists, and have been demonstrated to increase host virulence. The increasing ease of bacterial genome sequencing provides an opportunity to deeply explore prophage prevalence and insertion sites. Here we present VIBES, a workflow intended to automate prophage annotation in complete bacterial genome sequences. VIBES provides additional context to prophage annotations by annotating bacterial genes and viral proteins in user-provided bacterial and viral genomes. The VIBES pipeline is implemented as a Nextflow-driven workflow, providing a simple, unified interface for execution on local, cluster, and cloud computing environments. For each step of the pipeline, a container including all necessary software dependencies is provided. VIBES produces results in simple tab separated format and generates intuitive and interactive visualizations for data exploration. Despite VIBES’ primary emphasis on prophage annotation, its generic alignment-based design allows it to be deployed as a general-purpose sequence similarity search manager. We demonstrate the utility of the VIBES prophage annotation workflow by searching for 178 Pf phage genomes across 1,072 *Pseudomonas* spp. genomes. VIBES software is available at https://github.com/TravisWheelerLab/VIBES.

## INTRODUCTION

Bacteriophages (phages), viruses that infect bacteria, are as ubiquitous as their hosts. They are found everywhere that we find populations of bacteria, from forest soils and the oceans to hydrothermal springs and the human gut. Phages pose a significant threat to bacteria: in marine ecosystems, up to one third of bacteria are killed by phages every day (1). The strong pressure exerted by the threat of phage infection has led bacteria to evolve antiphage defense systems. Restriction-modification systems (2) and CRISPR-Cas systems (3) rely on sensing and degrading invading nucleic acids and are among the most widespread antiphage systems; however, numerous additional antiphage defense systems utilizing diverse mechanisms have recently been discovered (4). Likewise, phages have evolved to become adept manipulators of host metabolism, allowing them to evade host defenses or improve conditions for viral replication (5).

Phages can be purely parasitic (lytic) and replicate at the expense of their bacterial hosts. However, in addition to lytic replication, temperate phages can alternatively undergo lysogenic replication in which the phage genome integrates into the host chromosome as a prophage. Temperate phages are common: approximately half of sequenced bacterial genomes contain at least one prophage (6). Prophages are replicated each time the host cell divides. Consequentially, prophages benefit from thriving hosts, which can incentivize the development of mutualistic phage-host relationships. Many prophages encode factors that benefit their hosts (7). For example, some phages carry genes that promote resistance to infection from competing viruses (8), while other phages encode virulence factors, aiding host pathogenicity (9).

Many bacterial species are lysogenized by filamentous phages in the Inoviridae family (10). The opportunistic pathogen *Pseudomonas aeruginosa* is frequently lysogenized by an Inovirus called Pf (11–13). Pf prophages maintain lysogeny by repressing transcription of their excisionase (14). In response to oxidative stress (15), nutrient limitation (16), or other factors (17), the Pf prophage excises from the chromosome and Pf virion replication is initiated. Pf virion replication plays a role in biofilm development by lysing cells in the center of a colony, releasing DNA that adds to biofilm structural integrity (18). Pf virions themselves also act as structural components in *P. aeruginosa* biofilms, protecting bacteria from desiccation and antibiotics (19, 20). Indeed, the presence of Pf virions in the airways of cystic fibrosis patients is associated with antibiotic resistance (21). Additionally, Pf virions are immunomodulatory and induce maladaptive antiviral immune responses that promote infection initiation (22) and interfere with wound healing by inhibiting keratinocyte migration (23).

The important and varied roles of prophages in bacterial communities — parasites, mutualists, and sometimes pathogenicity aides — motivates the development of high-quality software methods that identify and classify their integrations into bacterial genomes. Generally, phage sequence annotation tools are designed to annotate either prophage in whole bacterial genome sequence (WGS) or phage genome fragments in metagenomic datasets. Most recent approaches focus on annotation in a metagenomic context, as metagenomic datasets have proven to be rich sources of previously unknown viral sequences (10, 24). Viral annotation, particularly in metagenomic contexts, requires tools to strike a balance between sensitivity and speed. High sensitivity is necessary to overcome high mutation rates in viral proteins combined with an increased risk of sequencing error stemming from the low abundance of viral sequences in most metagenomic datasets. Meanwhile, reasonable labeling speed is required when annotating large datasets. As a result of these constraints, viral annotation software has generally converged on a few techniques for identifying viral sequences: sequence similarity search against large databases of viral genomes (25–27), machine learning approaches based on statistical features such as *k*-mer frequencies, with a recent emphasis on neural networks (28–30), or some combination of both approaches (31–34). Viral annotation tools with a focus on prophage annotation in whole genome sequence (WGS) (DBSCAN-SWA (26), PHASTER (25), Prophage Hunter (33)) typically include visual summaries that increase the legibility of their output. Of these, only DBSCAN-SWA is also available as command line software, leaving it the only bacterial WGS annotation software that can be run by researchers seeking to conduct large-scale analysis in a cluster or cloud computing environment.

Here, we introduce and describe VIBES, an automated command line workflow for annotation of bacterial WGS that emphasizes identification of prophage integrations. VIBES supplements standard bacterial gene labeling with in-depth analysis of prophage integrations, producing machine- and human-readable text output files coupled with interactive HTML visualizations that facilitate further analysis of output data. VIBES is designed to:

- annotate prophages with high sensitivity;
- annotate bacterial genes on input genomes, using Prokka (35);
- annotate viral genes within input viral genomes, using the PHROG database (36);
- accept a potentially large number of bacterial genomes and candidate phage genomes as input;
- automatically manage distribution of workload to cluster or cloud resources; and

create interactive HTML visuals that display the above, as well as displaying which regions mapping to user input viral genomes are most prevalent among all input bacterial genomes. VIBES provides pre-built, Apptainer-compatible Docker containers that minimize dependency management. VIBES is implemented as a Nextflow-driven workflow that automatically orchestrates hundreds of jobs in a cluster or cloud environment. We demonstrate this with an analysis searching for 178 Pf phage variants in 1,072 complete *Pseudomonas* spp. genomes. Notably, while VIBES is intended for use as a prophage, bacterial gene, and viral gene annotator, it is effectively a sequence annotation workflow manager and can be used to orchestrate large-scale search for non-viral sequences in non-prokaryotic genomes (see Discussion).

## METHODS

### Implementation

VIBES (**V**iral **I**ntegrations in **B**acterial genom**ES**) is an automated prophage search workflow that uses containerized components coordinated by the Nextflow workflow management software (37) to produce output tab-separated value (TSV) annotation tables accompanied by interactive HTML files that summarize matches to prophage sequences. To annotate prophage integrations, VIBES is provided with an input FASTA file containing all prophage sequences to seek and a collection of bacterial genomes in FASTA format to annotate with prophages; it performs search using the software *nhmmer* (38). To annotate bacterial protein-coding genes, rRNA, and tRNA, VIBES uses Prokka (35). To annotate query prophage protein-coding genes, VIBES uses the BATH protein-coding DNA annotation software (39) and the PHROG v4 prokaryotic viral protein database (36) by default. The user can optionally substitute their own viral protein database. Figure 1 provides a visual representation of the three independent annotation workflows, run in parallel and managed by VIBES, that produce bacterial gene annotations, viral sequence integrations, and viral gene annotations. Table 1 lists the tools and databases used within VIBES.

**Table 1:** Automatically Managed Dependencies.

| Tool | Purpose |
| --- | --- |
| BATH (39) | Annotation of protein-coding DNA on query prophage genomes |
| HMMER (*nhmmer*) (38) | DNA-to-DNA alignment to identify prophage integrations on bacterial genomes |
| PHROG v4 (36) | Viral gene annotation database |
| Prokka (35) | Bacterial genome annotation |

**Figure 1:**
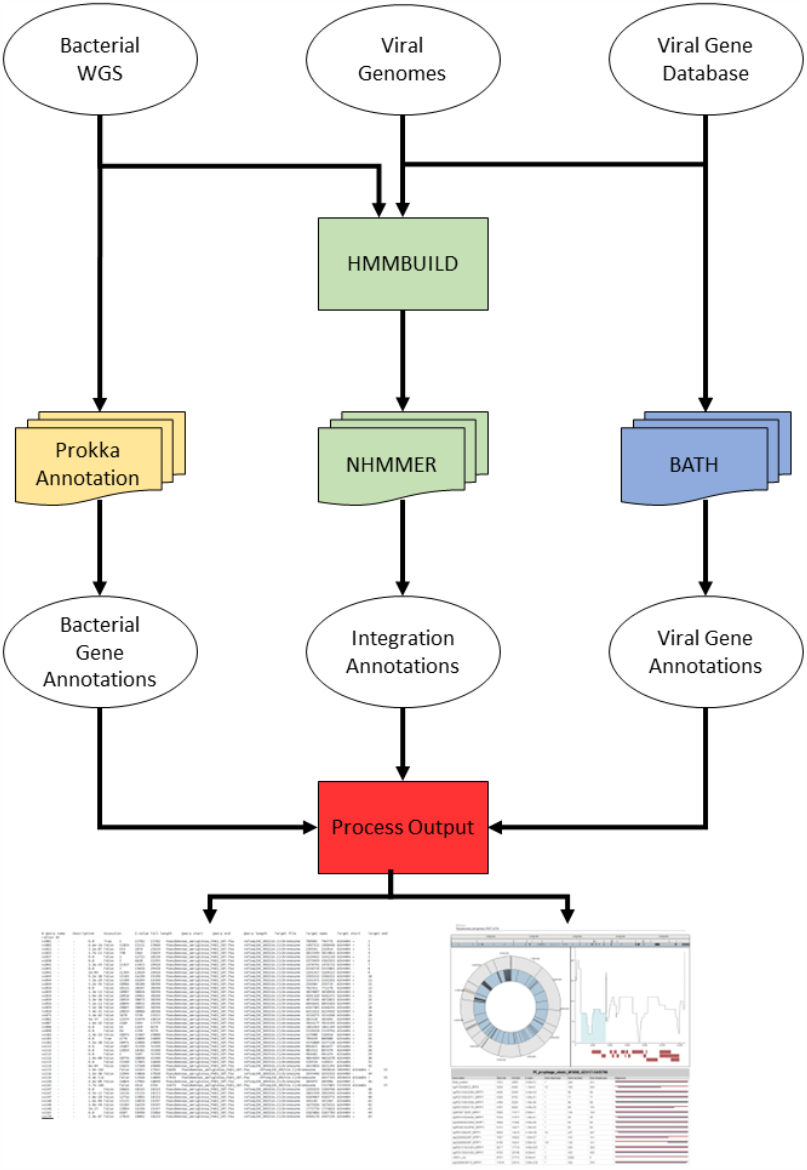
VIBES Workflow Schematic. This schematic displays how input data moves through the VIBES annotation workflow. Bacterial gene annotation processes are yellow, prophage annotation processes are green, viral gene annotation processes are blue, and visualization processes are red. The annotation processes are independent of each other. Stacked icons indicate processes parallelized automatically by Nextflow.

Before running VIBES, the user must install a software container system such as Docker (40) or Singularity/Apptainer (41) (usually the latter on HPC systems, where VIBES is likely to be utilized). These container systems enable the development and release of portable and reproducible software environments with fine-grained control over configuration and dependency conflicts while also retaining high performance. The user must also install the workflow management software Nextflow. Nextflow manages downloading and running containers, submits jobs to compute cluster job scheduling software (i.e. SLURM) or cloud computing architectures (i.e. AWS Batch), caches and checkpoints in-progress jobs in case of a crash, provides interpretable workflow status updates, runs on a wide variety of operating systems and hardware configurations, and can run VIBES locally as needed. After a user configures the workflow to run on their system and launches it, Nextflow requires no further user interaction to identify task dependencies, automatically maximizing parallelism by running as many tasks with satisfied dependencies as available resources allow.

The VIBES release consists of a Nextflow workflow script, several helper scripts written in Python and Perl, JavaScript and HTML files that produce the visualizations, and a Docker image that manages the internal configuration and dependency map of multiple tools. VIBES software and workflow can be found at https://github.com/TravisWheelerLab/VIBES.

### VIBES Components

#### Prophage Search Component

The primary component of the VIBES workflow is its prophage search. This component searches for user-provided query prophage sequences within bacterial genomes to identify prophage integrations. Identification of prophage within bacterial genomes is performed using a DNA sequence annotation tool, *nhmmer* (38), with default settings. Though it is slower than *blastn* (42), *nhmmer’s* improved sensitivity in the face of high sequence divergence and neutral mutation (38) is useful in the context of prophages, which can mutate at rates comparable to ssRNA viruses (43) and may show substantial divergence from query sequences. In general, any matches to a query prophage that fail to meet an E-value threshold (1e-5 by default) are discarded.

Sequence annotation tools such as *nhmmer* frequently produce fragmented alignments when presented with sequences highly diverged from the query sequences, particularly when a match contains a large inserted or deleted element relative to its nearest query. As a result, single prophage integrations may be reported as several fragments that lie close to each other on a bacterial genome in *nhmmer* output. To address these potentially fragmented integrations, VIBES includes a post-processing step that examines every match detected on a single bacterial genome, looking for consecutive matches that satisfy all of the following potential fragments must match to the same query phage sequence (Fig 2A), occur in the same order on the bacterial genome as on the query (Fig 2B), be close to each other on the bacterial genome (Fig 2Ca), and overlap minimally on the query phage (Fig 2Cb). Gaps between two matches on the query prophage sequence are not penalized, as they may represent large deletions. Matches that meet these criteria are assigned a common integration ID that instructs the interactive visual component to display the fragments together rather than separately (see Interactive Visuals), effectively joining the fragments into a single integration.

**Figure 2:**
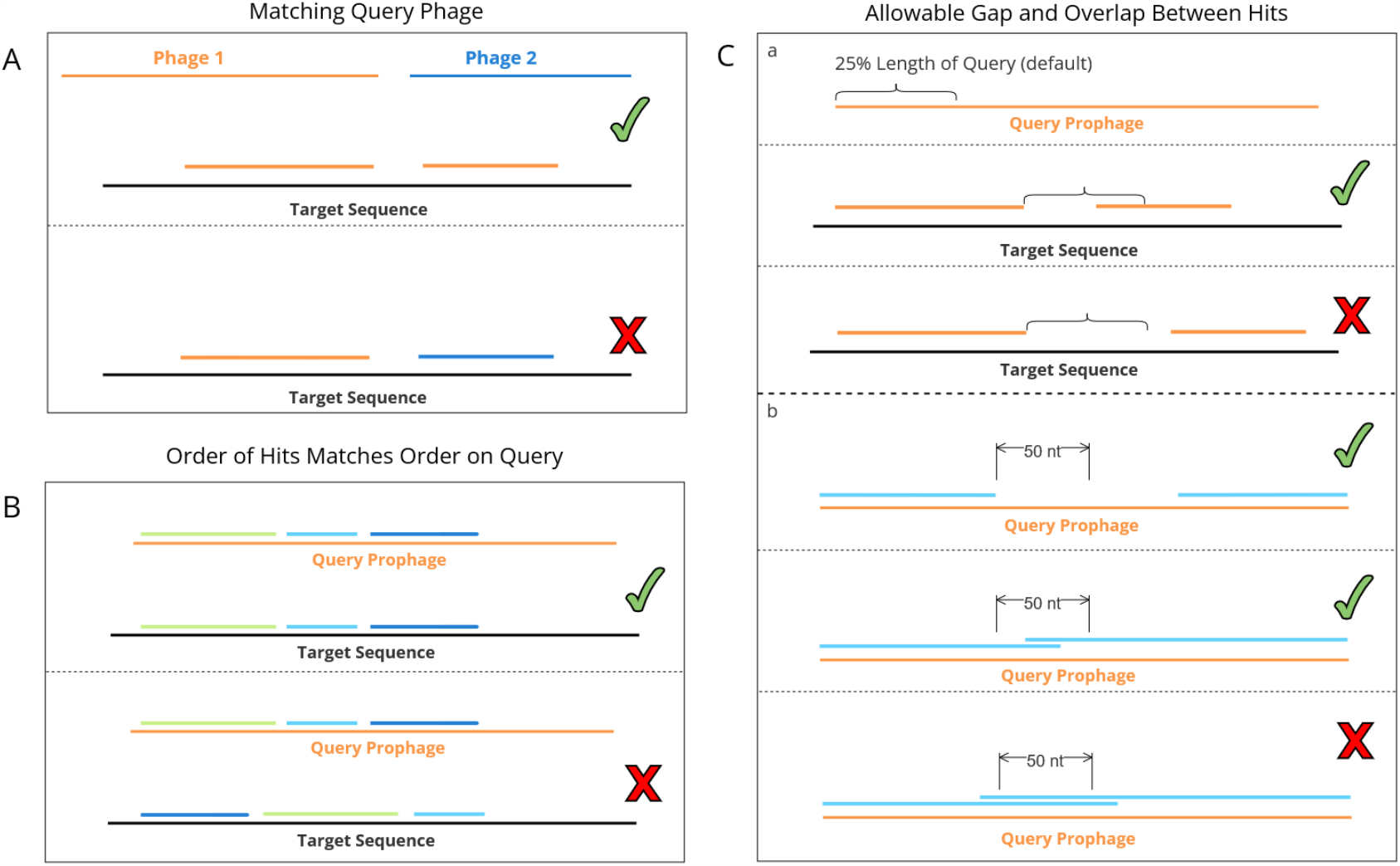
Diagram of Joining Parameters. Depicts all conditions that must be met for consecutive matches to be joined (assigned the same integration ID and displayed as one integration in visual output). **A**: Join candidates must be assigned to the same query phage. **B**: Join candidates must occur in the same order on both the query phage and target bacterial genomes. **Ca**: Given a query phage genome of length n, a match that ends at position i on the bacterial genome, and a consecutive match that begins at position j, two matches are considered near enough for fragment joining if |i − j| ≤ n ∗k, where k is a fragment gap threshold value set to 0.25 by default. **Cb**: Given a fragment whose match to the query viral genome ends at position s and a consecutive fragment whose match to the query viral genome begins at position t, the fragments are joined only when |t − s| ≤ θ, where θ is a constant set to 50 by default. Large gaps between matches on the query prophages are not penalized, as they may represent large deletions.

#### Gene Annotation of Both Viral and Bacterial Genomes

VIBES provides supplementary context to identification and investigation of prophage integration sites by identifying protein-coding genes in both full bacterial genomes and query prophage sequences. Each bacterial genome is annotated using the annotation tool Prokka (35) via StaPH-B’s Docker image (44), supporting gene annotation without requiring users to download or set up sequence databases. Like the prophage search component, each bacterial genome is annotated independently of other genomes, allowing Nextflow to fan out as many parallel Prokka annotation tasks as resources permit.

VIBES also produces gene annotations for the user-provided prophage sequences with its viral protein-coding gene annotation component. This component uses a translated search tool, BATH (39), to search a viral protein database against prophage DNA sequences. Translated search tools like BATH do not penalize neutral mutations that change DNA sequences without modifying the encoded protein sequence, making them especially well-suited to annotating sequences with high mutation rates such as viral genomes. BATH’s translated search is also robust to frameshift-inducing insertions or deletions, which can confound other translated search tools. By default, VIBES uses the PHROG v4 viral gene database (36) reformatted as a BATH-compatible HMM database, but users can substitute other amino acid sequence or HMM databases as desired.

Although VIBES was developed with annotating prophage integrations in mind, it is primarily a framework for managing and parallelizing runs of *nhmmer*, Prokka, and BATH with some prophage-annotation-specific features (the default PHROG database is phage-specific and the VIBES-SODA visualization suite assumes query sequences are prophage). In particular, the Prophage Search Component simply searches for matches to a query database (prophages by design) in a set of target genomes (bacteria by design) and can easily be repurposed by providing the workflow with a non-phage query sequence file and a set of non-prokaryotic genomes. Likewise, the Prokka bacterial gene annotation and BATH translated amino annotation components can be used to orchestrate massively parallel protein-coding sequence annotation, even on datasets where prophage integrations are not of interest.

#### Interactive Visual Generation

To facilitate further analysis and improve human readability of results, VIBES produces dynamic annotation visualizations in HTML files that can be opened in a web browser. These visuals depict prophage annotations, bacterial gene annotations, and viral gene annotations. After all other workflow tasks are complete, VIBES generates a collection of HTML files, each of which contains a dynamic visualization built with the SODA sequence annotation visualization library (45). The visualizations provide an interactive representation of VIBES output, including prophage annotations, bacterial gene annotations, and viral gene annotations. An output HTML file is generated for each input bacterial genome, each of which contains interactive annotation visualizations for its associated genome. The HTML files may be opened locally in a web browser, or they may be hosted on a web server. The generated interactive visualizations are described in Results.

### Pf Prophage Search

To demonstrate the utility of VIBES as a prophage identification tool, we searched 1072 *Pseudomonas* spp. isolate genomes for integrations of 178 Pf phage variants. *Pseudomonas* spp. genomes were acquired from the Pseudomonas Genome Database (v21.1) (46). Some records in the database were renamed to resolve characters that conflict with standard Bash commands while three records contained no sequence information. Two of the three empty records were populated with data from GenBank while the third was determined to be redundant and deleted (see Supplementary Data for details on modifications to data acquired from the Pseudomonas Genome Database). Phage sequence coordinates for 179 partial or complete Pf prophages were obtained from a study examining Pf prophage lineages (12). 126 *P. aeruginosa* genomes were downloaded using accession IDs provided in the study and the relevant sequences were extracted and assigned identifiers (see Supplementary Data for details). One phage sequence, labeled vs015, contained a substantial insertion that extended the length of the sequence to over 70kbp. Such a long query sequence requires a prohibitive amount of memory to search for, so vs015 was removed from our prophage database, leaving a total of 178 Pf phage query sequences derived from 126 *P. aeruginosa* genomes.

Analysis was conducted on the University of Arizona’s Puma HPC cluster on nodes that each contain 94 AMD EPYC 7642 cores and 512 GB of RAM.

## RESULTS

### Overview of VIBES Output Features

#### Detected Pf Prophages in Bacterial Genomes

For each input bacterial genome, VIBES produces a tab-separated value (TSV) file describing each detected potential prophage sequence in the genome. The TSV fields include matching phage name, match E-value, score, match start and end positions on both query (phage) and target (bacterial) sequences, match strand, a match integration ID (see Prophage Search Component under Methods and Materials), and a full-length field populated with True (full length) or False (partial). By default, a match is called full length if it is at least 70% the length of the best-matching prophage sequence, though this parameter can be modified by the user.

#### Bacterial and Viral Gene Annotation

Annotation of genes within bacterial genomes are generated by Prokka with its default annotation databases and settings. For each bacterial genome, full Prokka output is saved and optionally compressed into a zipped tar archive. Annotations of genes within prophage genomes are output in their own TSV format files with fields identical to those produced for prophage annotations except the match ID field, which is excluded for phage gene annotations.

#### Interactive HTML Visual Output

After each workflow process has completed, VIBES produces the SODA-based HTML visualization files. The interactive representations of the workflow’s output allows users to investigate annotations in a bacterial genome and potential prophages with the following components:

##### Bacterial Chromosome Plots (Fig 3B)

The visualizations include both circular and linear representations of the bacterial chromosome. Both representations of the chromosome are marked with detected viral integrations (yellow) and bacterial genes (blue) to assist in analysis of integrations and phage landing sites. Hovering over a blue bacterial gene marker displays the name of the gene, while hovering over a yellow phage integration marker displays the name of the prophage. Users can select a viral integration to inspect it more closely (see Position Occurrence Plot below). Users can zoom in on the chromosome and click and drag to pan across the genome, making gene or integration annotations larger and easier to interact with; simultaneously, the currently visible portion of the genome is highlighted in gray across the chromosome along top of the page. The circular genome can be changed to a linear representation, and vice versa, by clicking the linear button below the interactive chromosome.

**Figure 3:**
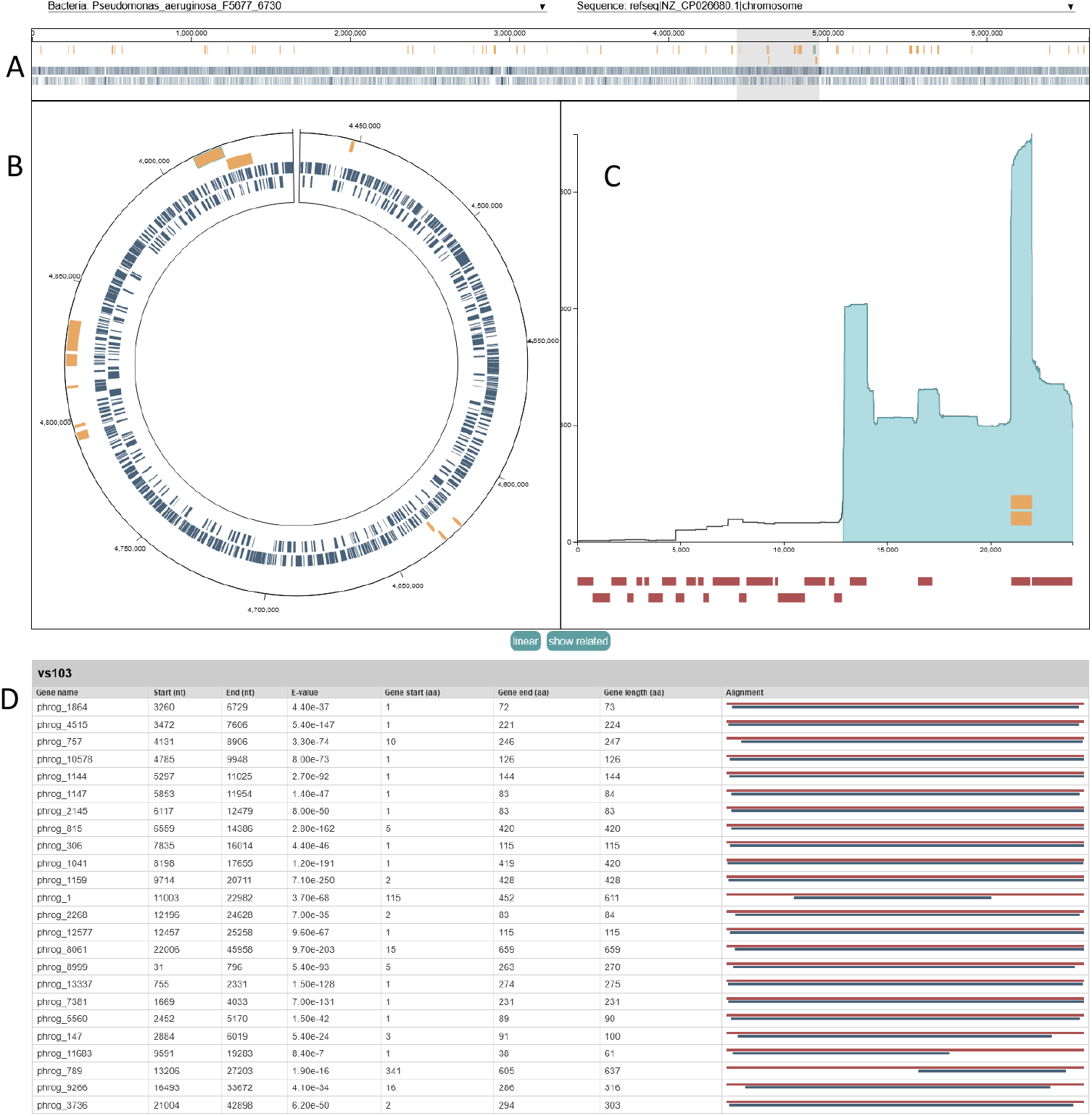
VIBES Interactive HTML Output. Example VIBES interactive annotation visualization page, displaying the bacterial chromosome with gene and prophage annotations, where the selected integration falls on the closest-matching viral genome, and viral gene annotations. **3A:** The full interactive visualization page. **3B:** The bacterial chromosome plot includes 2 modes to represent a bacterial chromosome: linear and circular, both marked with integration and gene annotations. **3C:** The position occurrence plot displays information about a selected integration, related integrations, and prophage gene annotations. **3D:** The query phage gene annotation table contains detailed information about gene annotations on the closest matching user provided phage genome.

##### Position Occurrence Plot (Fig 3C)

To assist users in investigating patterns of phage integration, a position-specific occurrence plot is displayed for a selected integration. The selected integration may be changed by clicking on a corresponding glyph in the genome annotation chart, or via the drop-down input at the top of the plot. The x-axis of the plot corresponds to each position (nucleotide) in a query phage sequence while the y-axis displays a count at each position summing every occurrence of that position in every integration in the dataset, emphasizing regions of a phage sequence that most often integrate into host genomes. The blue shaded region along the x-axis displays the extent of the currently selected integration on the query sequence it matched to. Yellow bars over the x-axis show where any other integrations matching to the same query phage on the selected bacterial genome matched to the query, indicating regions of the phage integrated repeatedly into the same genome. The yellow bars indicating where other integrations of the same phage fall on the viral genome can be hidden by clicking the hide related button located under the position occurrence plot. Under the x-axis, red bars display where viral gene annotations fall on the phage genome. Hovering over a viral gene annotation bar shows the name of the gene, while clicking on it highlights its row on the query phage gene annotation table.

##### Query Phage Gene Annotation Table (Fig 3D)

At the bottom of the visualization is a table of viral protein-coding gene annotations on query phage sequence most closely matching the currently selected integration. The phage gene annotation table contains a row for each annotated gene displaying the gene name, start and end positions on the query phage genome, annotation e-value, start and end positions relative to the reference gene amino acid sequence, and an alignment figure that visually depicts the extent of the match on the query phage sequence (blue line) compared to the reference amino acid sequence (red line).

### Application To *Pseudomonas* Dataset

To explore the utility of the VIBES workflow for identifying (possibly fragmented) phage integrations within bacterial isolates, we applied the workflow to a dataset is composed of 1,072 publicly available *Pseudomonas* spp. genomes obtained from the Pseudomonas Genome Database (v21.1) (46) and 178 Pf phage variants published in a study on Pf phage lineages (12).

Nextflow reported that the prophage detection component of the workflow consumed 13,526.3 CPU hours across 2,099 tasks in its prophage search component, 398.8 CPU hours across 1,072 tasks in its bacterial gene annotation component, and 49.7 CPU hours across 539 tasks in its viral gene annotation component, totaling 13,974.8 CPU hours consumed across a total of 3,710 tasks (more details on resource usage can be found in Table 2).

**Table 2:** Resource Usage.

| Component | Total CPU Hours | Tasks Run | Most Expensive Process (MEP) | CPUs Used, MEP | Mean RAM usage, MEP | Mean time, MEP (minutes) |
| --- | --- | --- | --- | --- | --- | --- |
| Prophage Search | 13,526.3 | 2,099 | *nhmmer* | 2 | 36.8 GB | 412.5 |
| Bacterial Gene Annotation | 398.8 | 1,072 | Prokka | 6 | 901.2 MB | 3.5 |
| Viral Gene Annotation | 49.7 | 539 | BATH | 2 | 403.5 MB | 8.6 |

In our input dataset of 1,072 target *Pseudomonas* spp. genomes and 178 query Pf phage variants, VIBES reported 51,386 partial and 517 full-length Pf phage integrations. Of the 51,903 integrations identified, 1,398 were composite integrations formed by 2 or more fragments joined together. The vast majority of reported integrations were less than 1,500 nucleotides in length (Fig 4). Although the workflow was set to discard matches less than 1,000 nucleotides in length, the median and average lengths of identified integrations were 1,240 and 2,419 respectively.

**Figure 4:**
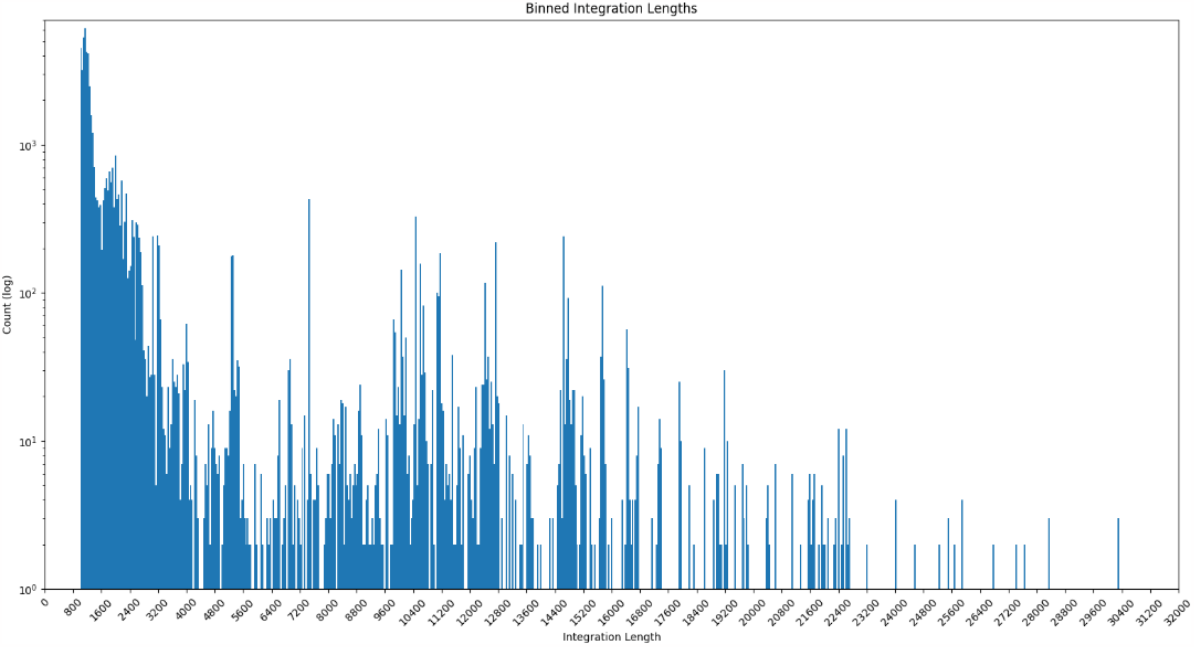
Integration Length Bin Plot. Counts of integrations binned by length, where joined integrations were summed together. The y-axis uses a log scale due to the large number of short integrations identified.

## DISCUSSION

Here, we have introduced a new software package that conveniently manages the workflow of annotating an arbitrarily large number of bacterial genomes, with special emphasis on identification of prophages. The VIBES workflow produces output annotation TSVs that are easy to load into a spreadsheet viewer or parse programmatically, along with helpful interactive HTML visualizations that facilitate analysis of identified prophages, their landing sites, and protein-coding genes on query phage sequences. VIBES also provides a basic framework for the management of general-purpose, large-scale annotation projects.

### Potential for Fine-Grained Prophage Search

Rather than *de novo* search for unannotated prophages, the relative sensitivity of the VIBES workflow’s prophage search may be useful in fine-grained searches assigning specific isolates to prophages flagged by a *de novo* annotation tool.

### Future Extensions

A common signal of phage integration is the presence of attachment (*att*) sites, two copies of which — *attP* and *attB*, carried on the phage genome and bacterial genome, respectively — flank integrated prophages. These leave a distinctive signal of flanking direct repeats, but this signal is not detected or annotated in the initial VIBES release. As *att* sites are of interest to prophage researchers and are annotated by some prophage annotation tools (26, 47), extending the VIBES feature set to include *att* site annotation is a high priority for future extensions of the workflow.

## AUTHOR CONTRIBUTIONS

C.J.C. developed the workflow software and performed all experiments. J.W.R. developed the visualization software component. P.R.S. and A.K.S. guided prophage analysis requirements and data acquisition. T.J.W. designed the approach and guided all research efforts. C.J.C and T.J.W. wrote the paper. All authors edited and approved the final manuscript.

## ACKNOWLEDGEMENTS

We are grateful to George Lesica for his early guidance in building Nextflow workflows. We also thank Elizabth Burgener, Julie Portois, and Paul Bollyky for sharing their data and analysis for use in evaluating VIBES during development. Analyses were made possible thanks to high performance computing (HPC) resources supported by the University of Arizona TRIF, UITS, and Research, Innovation, and Impact (RII) and maintained by the UArizona Research Technologies department.

